# Stem cell proliferation and differentiation during larval metamorphosis of the model tapeworm *Hymenolepis microstoma*

**DOI:** 10.1101/2023.09.13.557592

**Authors:** Jimena Montagne, Matías Preza, Uriel Koziol

**Affiliations:** Sección Biología Celular, Facultad de Ciencias, Universidad de la República, Montevideo, Uruguay

**Keywords:** differentiation, stem cell, neoblast, cestode, oncosphere, metacestode, tegument, neuroD

## Abstract

Tapeworm larvae cause important diseases in humans and domestic animals. During infection, the first larval stage undergoes a metamorphosis where tissues are formed *de novo* from a population of stem cells called germinative cells. This process is difficult to study for human pathogens, as these larvae are infectious and difficult to maintain in the laboratory. In this work, we analyzed cell proliferation and differentiation during larval metamorphosis in the model tapeworm *Hymenolepis microstoma*, by *in vivo* labelling of proliferating cells with the thymidine analogue 5-ethynyl-2′-deoxyuridine (EdU), tracing their differentiation with a suite of specific molecular markers for different cell types. Proliferating cells are very abundant and fast-cycling during early metamorphosis: the total number of cells duplicates every ten hours, and the length of G2 is only 75 minutes. New tegumental, muscle and nerve cells differentiate from this pool of proliferating germinative cells, and these processes are very fast, as differentiation markers for neurons and muscle cells appear within 24 hours after exiting the cell cycle, and fusion of new cells to the tegumental syncytium can be detected after only 4 hours. Tegumental and muscle cells appear from early stages of metamorphosis (24 to 48 hours post-infection); in contrast, most markers for differentiating neurons appear later, and the detection of synapsin and neuropeptides correlates with scolex retraction. Finally, we identified populations of proliferating cells that express conserved genes associated with neuronal progenitors and precursors, suggesting the existence of tissue-specific lineages among germinative cells. These results provide for the first time a comprehensive view of the development of new tissues during tapeworm larval metamorphosis, providing a framework for similar studies in human and veterinary pathogens.

## Introduction

Tapeworms (cestodes) are a diverse group of parasitic flatworms, many of which cause important diseases in humans and domestic animals (Budke et al., 2009; Torgerson et al., 2010; Hotez et al., 2014). Tapeworms have a divergent morphology and development that results from their adaptation to the parasitic lifestyle, including complex life cycles with successive larval and adult stages inhabiting different hosts (Freeman, 1973; Koziol, 2017). The first larval stage of tapeworms, called the oncosphere, is a miniaturized organism, typically containing fewer than a hundred cells (Ubelaker, 1980; Swiderski et al., 2016). This larva is specialized for the infection of the first host of the life cycle, and contains structures such as penetration glands and larval hooks, actuated by a complex system of muscles, which participate in the penetration of the intestine of the host. The oncosphere undergoes a metamorphosis in a parenteral site of the intermediate host, developing into the next life stage, the metacestode. Metacestodes from different cestode groups have a bewildering diversity of morphologies, which may or not include protective cyst tissues, but in all cases the infective metacestode includes an anterior scolex (head) with attachment organs, which is infective to the definitive host (Freeman, 1973; Chervy, 2002). Tapeworm species that cause the most important and life-threatening diseases are those in which humans are infected by larval forms (Budke et al., 2009), and the larval metamorphosis is thought to be a key step of the infection process during which the parasite is most vulnerable (Nono et al., 2012).

It is thought that most of the differentiated cells of the oncosphere are discarded during the larval metamorphosis (Freeman, 1973; Koziol, 2017). The differentiated tissues of the metacestode are therefore generated *de novo*, including the nervous and excretory systems, or extensively remodelled, including the tegumental syncytium that covers the larva. In tapeworms, and more generally in flatworms, differentiated cells are invariably post-mitotic and cell proliferation during post-embryonic development depends on undifferentiated stem cells (Reuter and Kreshchenko, 2004; Egger et al., 2009; Rink, 2013; Koziol et al., 2014; Rozario et al., 2019). These are usually called germinative cells in tapeworms, and are equivalent in their function to the neoblasts of free-living flatworms, such as planarians. Recent studies have shown that germinative cells are heterogeneous in their gene expression patterns, indicating that they are not a single cell population, but may comprise different lineages and hierarchies, as has been shown for planarian neoblasts (Koziol et al., 2014; Rozario et al., 2019; Molina and Cebriá, 2021).

Oncospheres of many different tapeworm species have been shown to possess a limited number of set-aside germinative cells, from which all further development is thought to occur after infection (Ubelaker, 1980; Koziol, 2017). However, very little is known about cell proliferation and differentiation during the larval metamorphosis in tapeworms, and it is unclear how these processes may relate or differ to other life-stages in tapeworms, or to other flatworm species. Most of what is known of larval metamorphosis in tapeworms comes from classical histological and electron microscopy studies, showing massive accumulation of germinative cells during the early metamorphosis, tegumental remodelling, and in some cases also including snapshots of the development of muscle and nerve cells (Bilqees and Freeman, 1969; Collin, 1970; Sakamoto and Sugimura, 1970; Shield et al., 1973; Schramlova and Blazek, 1983, Bortoletti and Ferretti, 1985; Holcman et al., 1994; Korneva, 2004). The study of larval metamorphosis in tapeworms that are directly relevant to human health, such as the taeniid genera *Echinococcus* and *Taenia*, is very difficult due to their infectivity to humans, the requirement for vertebrate laboratory hosts, and their slow metamorphosis (which can take weeks or even months, depending on the species, to reach metacestode infectivity). Although *in vitro* culture systems that support larval metamorphosis have been developed for some tapeworm species, these rarely allow complete development until infectivity and show a high variability (Heath and Smyth, 1970; Evans, 1980; Chile et al., 2016; Palma et al., 2019).

Species of genus *Hymenolepis* have been some of the most important laboratory models for the study of tapeworm biology, since the maintenance of their life-cycle in the laboratory is simple, using beetles and rodents as intermediate and definitive hosts, respectively (Arai, 1980). Recent developments have renewed the potential of these model species, including high quality genome sequences, life-stage and region-specific transcriptomes, optimization of methods for *in situ* gene expression analysis, and functional analyses by RNA interference (Cunningham and Olson, 2010; Pouchkina-Stantcheva et al., 2013; Olson et al., 2018; Rozario et al., 2019; Olson et al., 2020; Preza et al., 2021). The metacestode of *Hymenolepis* spp. is called a cysticercoid, and once developed consists of a small scolex and body that are withdrawn within a protective cyst (also called capsule). A tail or appendage, called the cercomer, continues growing from the posterior of the cyst even after infectivity is reached. Once the cysticercoid is ingested by the definitive host, the cyst and cercomer tissues are destroyed in the digestive system, and the freed activated cysticercoid consists solely of the scolex and a small posterior body. Early studies described only in broad strokes the larval metamorphosis of different *Hymenolepis* species, mostly by classic histological methods (Ubelaker, 1980). More recently, a transcriptomic analysis of the early metamorphosis in *Hymenolepis microstoma* has shown similarities in global gene expression to the generative neck region of adult worms, and described the localized expression of several transcription factors indicating that larval metamorphosis is a highly dynamic process, in which fate specification and cell differentiation are likely to begin from the earliest stages (Olson et al., 2018). In this work, we have studied in detail the early larval metamorphosis of the model tapeworm *H. microstoma*, describing the development of the different tissues and systems of the metacestode, and tracing the proliferation and differentiation of germinative cells during these processes.

## Materials and methods

### Parasite material

*H. microstoma* (“Nottingham strain”) was maintained using C57BL/6 mice as definitive hosts, and *Tribolium confusum* as intermediate hosts, as previously described (Cunningham and Olson, 2010), in collaboration with Jenny Saldaña, Laboratorio de Experimentación Animal, Facultad de Química, Universidad de la República, Uruguay (“Mantenimiento del ciclo vital completo del cestodo *H. microstoma* utilizando sus hospedadores naturales Mus musculus (ratón) y *Tribolium confusum* (escarabajo de la harina)”, protocol number 10190000025215, approved by Comisión Honoraria de Experimentación Animal, Uruguay).

Starved beetles were exposed to infective eggs overnight, and routinely incubated at 28 ± 1 °C. For some infections, incubation was performed at lower (25 °C) or higher (30 °C) temperatures to accelerate or delay development in order to accommodate the times at which larvae at specific developmental stages had to be collected. *In vitro* activation of infective cysticercoids was performed as described by Preza et al., 2022. Larvae were routinely fixed for most downstream protocols using 4% paraformaldehyde prepared in phosphate buffered saline (PBS), overnight at 4-8 °C.

### Infection of *Tenebrio molitor* and *in vivo* labelling with 5-Ethynyl-2’-deoxyuridine (EdU)

*T, molitor* beetles were purchased from local providers, or raised at 20-26°C with a 12 h:12 h photoperiod, in 20 x 10 cm pots with 90% wholemeal flour and 10% yeast, plus fruit once a week. Beetles were starved for 48 hours before being exposed to infective eggs overnight. For EdU labelling, beetles were anaesthetised with triethylamine for one to two minutes, the posterior half of one elytron was removed with forceps, and 5 μl of a 200 μM solution of EdU (Thermo-Fisher) was injected with a Hamilton precision syringe into the hemocoel in the abdomen. Beetles were maintained in wholemeal flour at 28 ± 1 °C, and larvae were collected after different times by dissection of the beetles in PBS. EdU labelling was developed with the Click-iT™ EdU Cell Proliferation Kit for Imaging, Alexa Fluor™ 555 dye (Thermo-Fisher C10338).

### Dextran labelling of the tegument

Larvae were collected by dissecting infected beetles in PBS, and stained with tetramethylrhodamine and biotin conjugated dextran (10,000 MW, Lysine Fixable, Thermo-Fisher D3312) with a protocol modified from Wendt et al., 2018. Up to 300 larvae were incubated in 250 μl of a 2-5 mg/ml solution of conjugated dextran; larvae collected during the first two days post infection were incubated briefly without vortexing, whereas larvae collected from three days post-infection onwards were incubated for 2 to 3 min with low speed vortexing. Then, larvae were fixed by adding 1 ml of 4% paraformaldehyde prepared in PBS, and the solution was replaced immediately with 1 ml of fresh 4% paraformaldehyde solution and incubated overnight at 4-8 °C.

### Transmission Electron Microscopy (TEM)

Larvae collected three days post infection were fixed in glutaraldehyde (2.5 %) and paraformaldehyde (2%) prepared in cacodylate solution (50 mM cacodylate, 50 mM KCl, 2.5 mM MgClL pH 7.2) for 3 hours. After washing, samples were post-fixed in 1 % osmium tetroxide, dehydrated using ethanol, infiltrated and embedded in Araldite resin. Ultrathin 70 nm sections were obtained using an RMC MT-X ultramicrotome and mounted on formvar-coated copper grids. Observation and acquisition was performed using a Jeol JEM 1010 transmission electron microscope operated at 100 kV, equipped with a Hamamatsu C4742-95 digital camera (Unidad de Microscopía Electrónica, Facultad de Ciencias, Universidad de la República, Uruguay).

### Whole mount immunofluorescence (WMIHF)

WMIHF was performed following a modification of the protocol described by Koziol et al., 2013. Briefly, fixed larvae were washed extensively with PBS containing 0.3% Triton X-100 (PBS-T), permeabilized for 20 minutes in PBS containing 1% sodium dodecyl sulfate (SDS), washed again three times in PBS-T, and blocked for 2 hours in PBS-T with 3% bovine serum albumin (BSA, Merck) and 5% normal sheep serum (Merck). Incubation with primary antibodies was done for 12 to 72 hours at 8 °C in PBS-T with 3% BSA and 0.02% sodium azide. After four one-hour long washes, the samples were incubated with secondary antibodies for 12 to 72 hours at 8 °C in PBS-T with 3% BSA and 0.02% sodium azide. Finally, samples were washed four times for one hour in PBS-T. Nuclei were stained using either 4′,6-diamidino-2-phenylindole (DAPI) or methyl green (Prieto et al., 2015), and in some experiments actin filaments were stained with phalloidin conjugated to fluorescein isothiocyanate (Merck). The primary antibodies used were: rabbit polyclonal anti-tropomyosin (Koziol et al., 2011), 1:500 dilution; rabbit polyclonal anti-FMRFamide (Immunostar, ID 20091), 1:300 dilution; rabbit polyclonal anti-serotonin (Immunostar, ID 20080), 1:300 dilution; rabbit polyclonal anti-phospho-histone H3 (Ser10) (Cell Signalling Technology #9701) 1:100 dilution; mouse monoclonal anti-synapsin (clone 3C11, Developmental Studies Hybridoma Bank), 1:100 dilution. The secondary antibodies used included anti-rabbit antibodies conjugated to Alexa Fluor 546 (Invitrogen A11010) and anti-mouse antibodies conjugated to Alexa Fluor 647 (Invitrogen A31571).

### Histological sectioning and immunofluorescence

For some experiments, immunofluorescence was performed with 20 μm-thick cryosections of samples previously included in Tissue-Tek OCT compound (Sakura FineTek cat. no. 4583) as previously described (Koziol et al., 2013).

### Identification of *H. microstoma* homologs of transcription factors related to muscle and neural development

The predicted proteome of *H. microstoma* was downloaded from WormBase ParaSite (Howe et al., 2017; https://parasite.wormbase.org/, version 14), and used to search by reciprocal BLASTP for homologs of *myoD*, *nkx-1.1*, *coe* (collier), *neuroD* and *soxB*, using sequences from *Homo sapiens*, *Drosophila melanogaster*, *Caenorhabditis elegans*, and *Schmidtea mediterranea*. Further confirmation of the orthology of basic helix-loop helix genes (bHLH) was obtained by InterPro domain analysis (Paysan-Lafosse et al., 2023). In the case of *neuroD*, InterPro domain analysis confirmed the presence of the *neuroD-*specific domain IPR022575 at the C-terminus of the predicted protein. For *coe*, InterPro domain analysis also confirmed the presence of the specific COE DNA-binding domain (IPR032200) and IPT domain (IPR002909). For *myoD*, InterPro domain analysis confirmed the presence of the specific MyoD_N N-terminal domain (IPR002546).

### Whole mount in situ hybridization (WMISH)

Fragments of the coding sequence of each gene were obtained by RT-PCR of *H. microstoma* total adult cDNA using specific primers (Supplementary Table S1), cloned into pGEM-T (Promega) and used to synthesize digoxigenin-labeled probes by in vitro transcription using SP6 or T7 polymerases (Thermo-Fisher), in reactions containing 3.5 mM digoxigenin-UTP (Merck), 6.5 mM UTP, and 10 mM ATP, GTP and CTP. The cDNA fragments used for probe synthesis for *pc2*, *chat*, *vglut* and *tph* were the same as the ones described in Preza et al., 2018. Fluorescent whole-mount in situ hybridization (WMISH) was performed as previously described (Koziol et al., 2014; Koziol et al., 2016). Samples were co-stained with DAPI or methyl green. When WMISH was combined with EdU labelling or WMIHF, these protocols were performed after WMISH was complete.

### Sample mounting, imaging and image analysis

Samples were mounted in 80% glycerol with 50 mM Tris, pH 8.0, or with ProLong Glass Antifade Mountant (Thermo-Fisher). Samples were imaged by confocal microscopy (Zeiss LSM 800CyAn and Zeiss LSM 880, Advanced Bioimaging Unit of the Institute Pasteur of Montevideo). Images were analyzed and processed using FIJI (Schindelin et al., 2012).

### Statistics

Statistical analysis was carried out using Graphpad Prism 8 software for plots, linear regression and non parametric Mann-Whitney test (significance was considered at p < 0.05).

## Results

### Overview of larval metamorphosis in *H. microstoma*

An overview of the metamorphosis of *H. microstoma* was described by Voge (1964) and Goodchild and Stullken (1970). We have modified the staging system of Voge (1964), subdividing the developmental stages based on finer anatomical details. These stages, and their corresponding time of appearance (in days post infection, d.p.i.) at 28 °C, are shown in Figure 1, and described briefly below:

**Figure 1.**
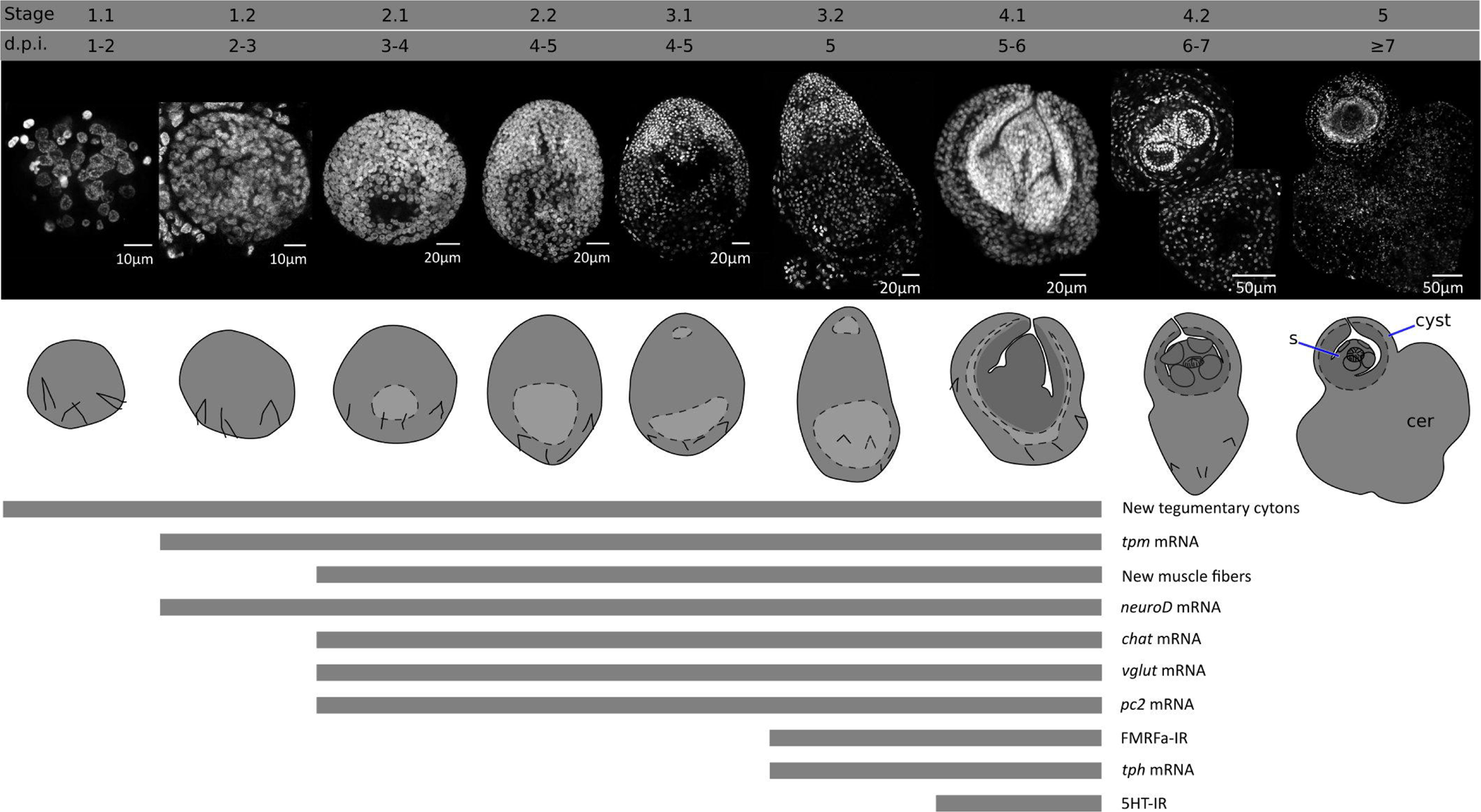
Developmental stages of larval metamorphosis in *H. microstoma*. The morphology at each developmental stage is shown by nuclear staining (top pictures) and diagrams (bottom pictures). The presence of some differentiation processes and markers studied in this work, up to stage 4.1, is summarized with bars below. Abbreviations: s, scolex; cer, cercomer.

Stage 0 (0 d.p.i). We refer to the hatched infective oncosphere as stage 0 (as this was not included in the staging system of Voge, 1964). Larval diameter is approximately 25 to 30 µm.

Stage 1.1 (1-2 d.p.i). The larva is compact, has a roughly circular outline, and has a diameter of approximately 35-50 µm.

Stage 1.2. (2-3 d.p.i.). The larva has a roughly circular outline, with a diameter of approximately 50-70 µm. A small central cavity begins to form between the deeper cells, this is the beginning of the formation of the so-called central cavity, also known as the primary lacuna (Freeman, 1973).

Stage 2.1 (3-4 d.p.i.). The outline of the larva is ovoid. The central cavity becomes larger, and is displaced to the posterior (closer to the remaining oncospheral hooks). Nuclei in the anterior region, which will become the scolex, are smaller and more compactly distributed. The length of the larva is 70-110 µm.

Stage 2.2 (4-5 d.p.i.). The outline of the larva is ovoid, but further growth occurs and the central cavity becomes larger. The length of the larva is 95-150 µm.

Stage 3.1 (4-5 d.p.i.). The outline of the larva is ovoid, and the primordia of the attachment organs of the scolex become distinguishable. The primordium of the rostellum (apical attachment organ) can be distinguished by a gap between the primordium and the rest of the larva, and the primordia of the four suckers can be seen as small masses of cells. The length of the larva is 150-200 µm.

Stage 3.2 (5 d.p.i). The larva becomes elongated, and the primordia of the suckers become more prominent. The length of the larva is 200-270 µm

Stage 4.1 (5-6 d.p.i.). The anterior part of the larva (the future scolex and body of the cysticercoid) withdraws into the cavity, becoming surrounded by the future cyst tissues. The cavity collapses as the anterior end withdraws. The withdrawal of the scolex and body into the cyst tissues has been shown to be a fast process mediated by the retraction of muscle fibers (Caley, 1974). The scolex primordium is thus located at the bottom of the withdrawn tissue, and the developing body is folded around the scolex. The posterior-most region of the larva begins to grow, becoming the cercomer (tail).

Stage 4.2 (6-7 d.p.i.). Scolex development proceeds, including the differentiation of the rostellum (as rostellar hooks appear) and suckers. The cyst tissues develop around the scolex and body. The cercomer grows to a size comparable to that of the cyst.

Stage 5 (8 d.p.i). Scolex and cyst development concludes. Cercomer growth may continue beyond this stage, and reach lengths that are many times longer than the cyst.

There is some variability in the developmental stages of larvae found between different beetles and within the same beetle, which may be due in part to the prolonged overnight exposure of the beetles to the infective eggs, resulting in slightly different infection times.

### Cell proliferation during larval metamorphosis

Growth is very fast during the early stages of metamorphosis, and the total number of nuclei increases exponentially during the first three days of metamorphosis (from an average of 43 nuclei 1 d.p.i., to 1219 nuclei 3 d. p. i., Figure 2A). The doubling time for the total number of nuclei was estimated to be 10 hours. Because cell cycle exit and differentiation are already underway for some cells at these stages (see below), this gives an upper bound to the total length of the cell cycle of proliferating cells during early larval metamorphosis.

**Figure 2.**
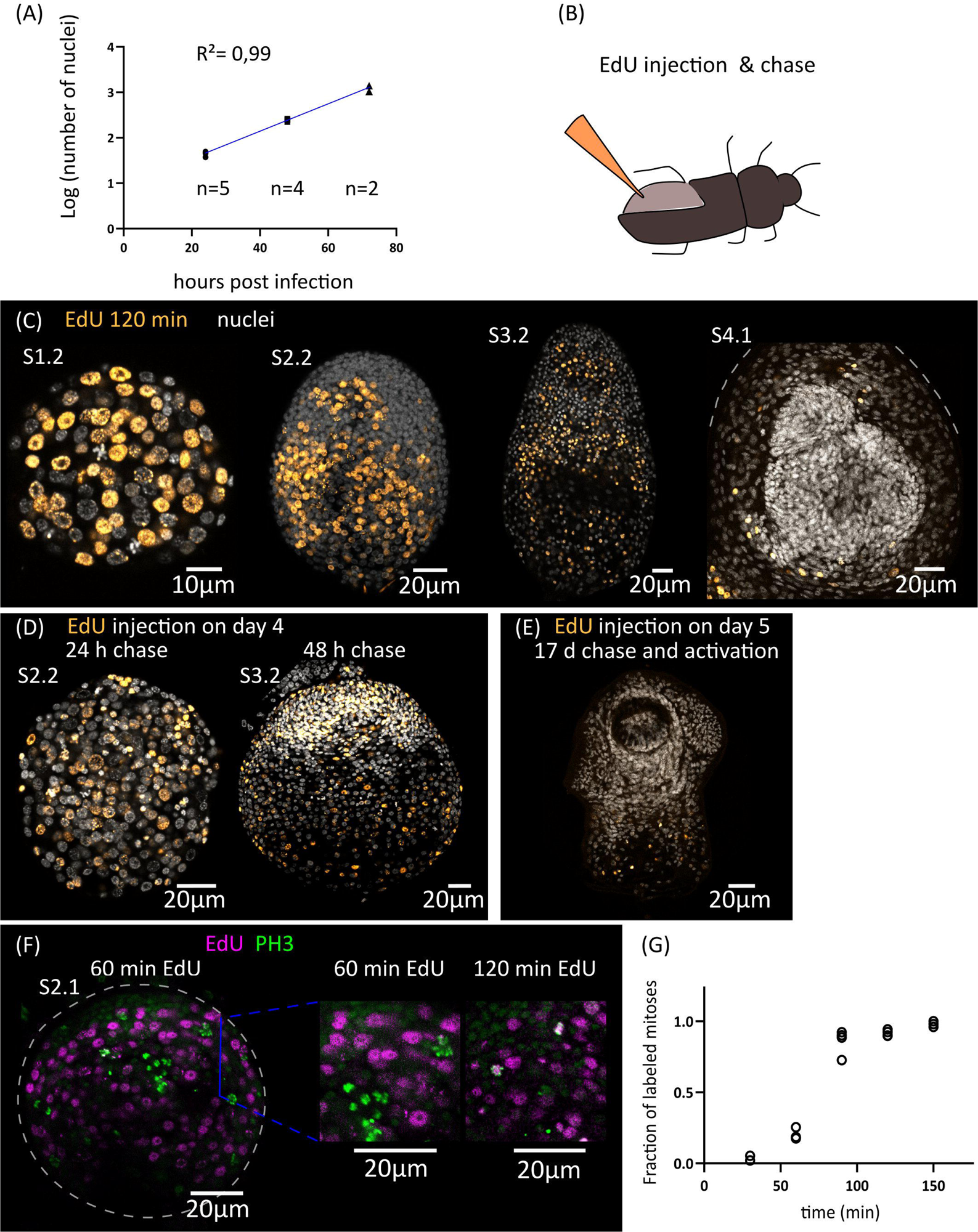
Cell proliferation during larval metamorphosis in *H. microstoma*. (**A).** Quantification of the increase in the total number of nuclei during early larval metamorphosis**. (B)** Diagram showing the method used for EdU injection in infected *T. molitor* beetles. **(C)** EdU labelling at 2 h after injection for larvae at different developmental stages. **(D)** EdU labelling in larvae at 24 to 48 h of chase after injection. **(E)** EdU labelling in a cysticercoid larva that was exposed to EdU at 5 d.p.i., allowed to fully develop *in vivo* for an additional 17 days, and activated by mimicking *in vitro* the infection of the definitive host. **(F)** Detection of EdU and phosphorylated histone H3 (PH3) in developing larvae at different time points after the injection of EdU to infected beetles. **(E)** Quantification of the fraction of labelled mitoses in larvae at different time points after the injection of EdU to infected beetles.

In order to identify the distribution of proliferating cells during larval metamorphosis, we developed an *in vivo* protocol for metabolic labelling of cells undergoing DNA synthesis using the thymidine analogue 5-ethynyl-2′-deoxyuridine (EdU). To this end, we infected *Tenebrio molitor*, a large beetle species susceptible to infection by *H. microstoma* (Voge and Graiwer, 1964) which can be easily manipulated experimentally. EdU was injected into the hemocoel of the beetle (Figure 2B) at different time points after infection, and the incorporation of EdU by *H. microstoma* larvae was assessed 2 hours after injection (Figure 2C). In stages 1.1 and 1.2, EdU^+^ cells are very abundant, and distributed throughout the larval tissues. Their nuclei are relatively large, and only a few smaller nuclei in the periphery of the larvae appear to always be EdU negative. In later stages (2.1 to 3.2), although EdU^+^ cells are still abundant, there are many regions that appear to be devoid of proliferation, especially at the surface of the larva and in specific regions of the scolex, including the anterior pole. Finally, in stages 4.1 and 4.2 (after the withdrawal of the anterior end), EdU^+^ cells are absent from the developing scolex, but present in the cysticercoid body and cercomer. These results suggest that although development is still underway in the scolex after its retraction, this is based on the differentiation of cells that are already post-mitotic.

By performing similar experiments, but varying the time at which parasites were collected after EdU injection (“chase”), we were able to trace the fate of proliferating cells at different points of development. When larvae were exposed to EdU at stages 1.1 to 1.2 and allowed to develop for 24 to 48 hours, the progeny of the cells incorporating EdU could be detected throughout the larval tissues, with a strong accumulation in the periphery and scolex of larvae at stages 3.1 and 3.2 (Figure 2D). In contrast, when larvae were exposed to EdU at stages 4.1 to 4.2, allowed to complete metamorphosis, and activated *in vitro* (mimicking the infection of the definitive host), labelled cells were restricted to the body and absent from the scolex. Therefore, there is a pronounced antero-posterior gradient in the development of the cysticercoid, and only cell proliferation during the first developmental stages (up to stage 3.2) contributes to the formation of the scolex.

Finally, by combining EdU labelling with immunofluorescence for phospho-histone H3 (PH3), a mitotic marker, we performed an analysis of the fraction of labelled mitosis (FLM, Shackney and Ritch 1987) in order to estimate some cell cycle parameters during early larval metamorphosis (Figures 2F and 2G). Larvae at stages 2.1 to 2.2 were exposed to EdU *in vivo*, and collected at 30 min intervals for double detection of EdU and PH3. As expected, after only 30 minutes there is almost no co-localization of both labels, as cells that incorporated EdU are expected to be still in S phase or in G2 phase. The fraction of EdU labelled mitoses increased steadily with time as the labelled cells reached M phase, and almost all mitoses were labelled after 120 min. The average length of G2 can be estimated as the time point at which 50% of the mitoses are labelled (Shackney and Ritch 1987), which was approximately 75 minutes. In a different time-course experiment with longer chase times, we found that only 14% of mitoses were labelled after 27 hours. These results indicate that the duration of the EdU pulse in the hemocoel must be relatively short (if the EdU pulse lasted for 27 hours, the fraction of labelled mitoses would not drop within this time frame).

### Development of the larval tegument

Parasitic flatworms, including tapeworms, trematodes and monogeneans, are not covered by a typical epidermis but instead possess a syncytial tegument (Tyler and Hooge, 2004). The tegument consists of a superficial band of cytoplasm (the distal cytoplasm) that is connected by thin cytoplasmic bridges to individual nucleated cell bodies (cytons) lying beneath the basal lamina. In the oncosphere of most tapeworms, including *Hymenolepis* spp., a single binucleated cyton is present (Ubelaker, 1980), and new cytons arise during metamorphosis from the fusion of the progeny of germinative cells to the distal tegument.

We adapted to *H. microstoma* larvae a method for labelling the tegument with fluorescently conjugated dextran, originally developed by Wendt et al., 2018 for the trematode *Schistosoma mansoni*. Several cytons could be identified at 1 d.p.i., but we were unable to localize the original binucleated cyton of the oncosphere, suggesting that it was already eliminated at this time point. The number of cytons connected to the distal tegument increased proportionally to the total number of cells of the larvae during the first three days of metamorphosis, and were distributed throughout the surface of the larvae (Figure 3A, B). We confirmed the presence of tegumental cytons at stages 1.1 to 1.2 (3 d.p.i.) by transmission electron microscopy, with several short cytoplasmic processes connecting them to a very thin distal cytoplasm (approximately 200 nm in thickness) (Figure 3C).

**Figure 3.**
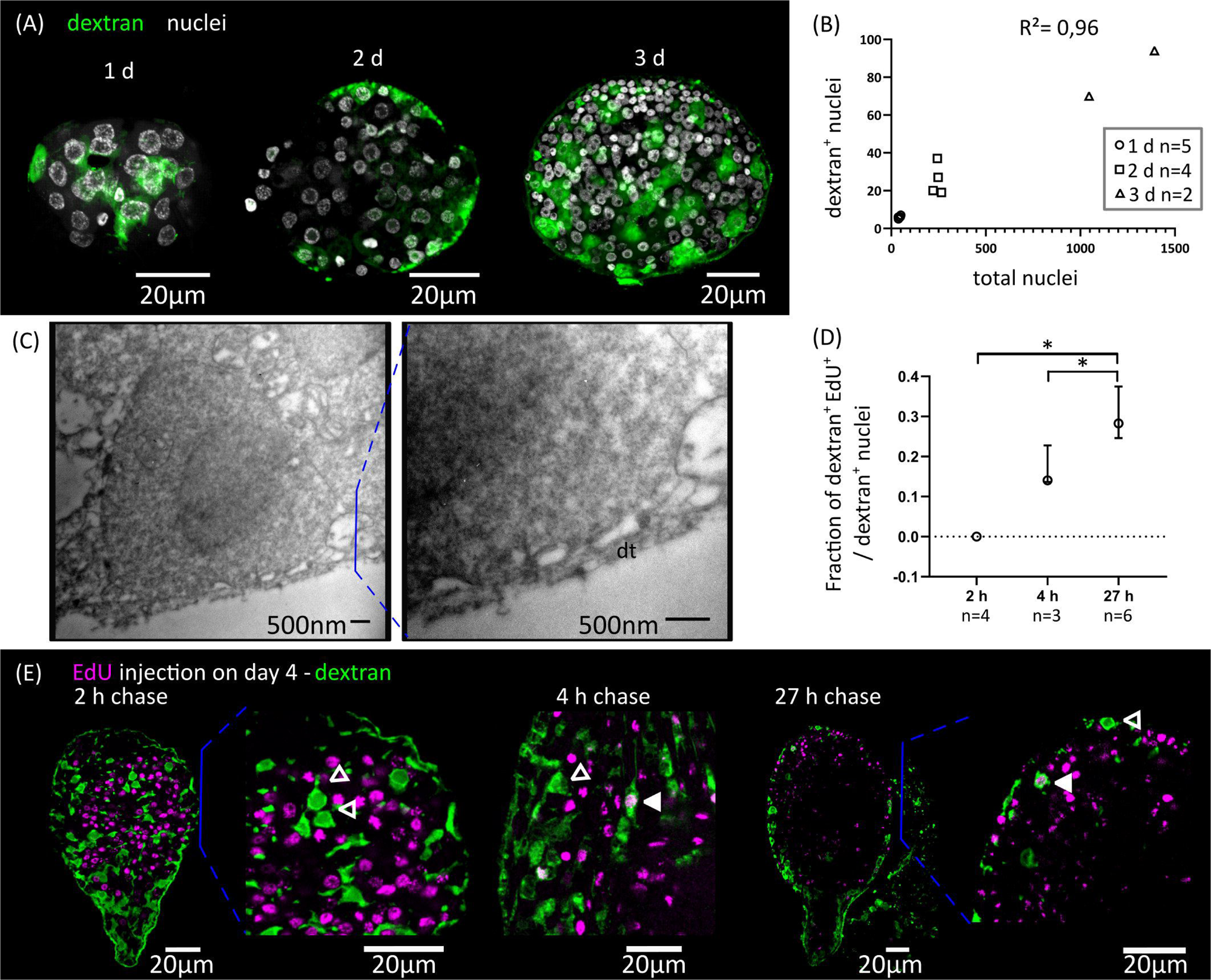
Differentiation of tegumentary cytons. **(A)** Dextran labelling of the tegument of larvae at different time points after infection (1, 2 and 3 d.p.i., corresponding to stages 1.1, 1.2 and 2.1). **(B)** Quantification of the increase of dextran-labelled nuclei in comparison to the total number of nuclei. **(C)** Transmission electron microscopy of a tegumentary cyton in a larva at 3 d.p.i. (left), including a detailed view of the distal tegument (dt) and its connection to the tegumentary cyton via cytoplasmatic bridges. **(D)** Quantification of the fraction of tegumentary cytons (dextran^+^ nuclei) that are labelled by EdU at different chase times after the injection of EdU to infected beetles. The points show median values and the whiskers show the maximum and minimum values (* p < 0.05, Mann-Whitney test). **(E)** Examples of larvae combining detection of EdU incorporation and dextran labelling of the tegument at different chase times after the injection of EdU to infected beetles. Open and filled arrowheads indicate dextran^+^/EdU^-^ and dextran^+^/EdU^+^ cytons, respectively.

We analyzed the kinetics of differentiation of germinative cells into tegumental cytons by combining EdU chase experiments with dextran labelling. We exposed larvae at stages 2.1 to 3.1 to EdU *in vivo*, and followed the fate of labelled cells (Figure 3D, E). Two hours after EdU exposure, all dextran^+^ tegumental cytons were negative for EdU, confirming that these are post-mitotic. However, EdU^+^ tegumental cytons could already be detected at 4 hours after EdU exposure (17% of all tegumental cytons were EdU^+^ in average), and the proportion of labelled cytons increased after 27 hours (29 % of all dextran^+^ tegumental cytons were EdU^+^ in average, Figure 3D).

In summary, fusion of new cells to the tegument starts at the very beginning of larval metamorphosis, and our results indicate that a constant proportion of the proliferative output of germinative cells is destined to this structure during the early metamorphosis. Fusion of precursors to the tegumental syncytium may be a very early step of the differentiation process, as it can be detected after only 4 hours of EdU labelling (less than three hours after the last mitosis, according to our estimate of the length of G2).

### Remodelling of the muscle system

The muscle system of the oncosphere comprises superficial muscle fibers (mostly circular fibers), a pair of dorso-ventral muscles, and a complex arrangement of muscles that are attached to the hooks and are responsible for their extension and retraction (Ubelaker, 1980; Hartenstein and Jones, 2003). This arrangement is replaced during larval metamorphosis, as the cysticercoid has a complex complement of muscular fibers that is most similar to the adult worm. This includes longitudinal and circular muscle fibers of the body wall (sub-tegumental muscle), inner longitudinal muscle fibers, transverse muscle fibers, and complex muscle arrangements in the attachment organs (rostellum and suckers).

We analyzed the remodelling of the muscle system by detecting muscle fibers using an antibody that recognizes muscular tropomyosin isoforms (high molecular weight tropomyosin isoforms: HMW-TPM; Koziol et al., 2011) (Figure 4A). Similar results were obtained by staining the larvae for actin filaments with fluorescent phalloidin (Figure 5). During the initial stages of metamorphosis (stages 1.1 to 1.2), the muscle fibers appear thin and discontinuous on the surface of the larvae, and the hook muscles appear globose and disorganized (Figure 4A). It is likely that these are the degenerating muscle fibers of the oncosphere. The first clear evidence of the differentiation of new muscle fibers appears at stage 2.1, in which circular muscle fibers and six bundles of longitudinal muscle fibers (two lateral bundles and two pairs of dorsal and ventral bundles) appear beneath the tegument of the body wall (Figure 4A). Surprisingly, although the muscle system of adult cestodes is dorsoventrally symmetrical, these early bundles of longitudinal muscle fibers show a dorsoventral asymmetry. On one side (due to the absence of other morphological landmarks, it is not possible to determine if this corresponds to the dorsal or ventral side), the bundles of fibers are thicker and more developed, and each bundle is clearly associated to a single large HMW-TPM^+^ cell body. At later stages (2.2 to 3.2), both circular and longitudinal muscle fibers increase their number and density beneath the tegument, forming an orthogonal grid (Figure 4A). The longitudinal fibers extend through most of the length of the larvae, but are absent in the posterior-most region that will become the cercomer, as described by Caley (1974). Additionally, transverse muscle fibers appear in the anterior region (the future scolex), particularly at the base of the developing rostellum, and between the sucker primordia. The intrinsic musculature of the rostellum and suckers only differentiates after scolex withdrawal, during stages 4.1 to 4.2 (Figure 4A).

**Figure 4.**
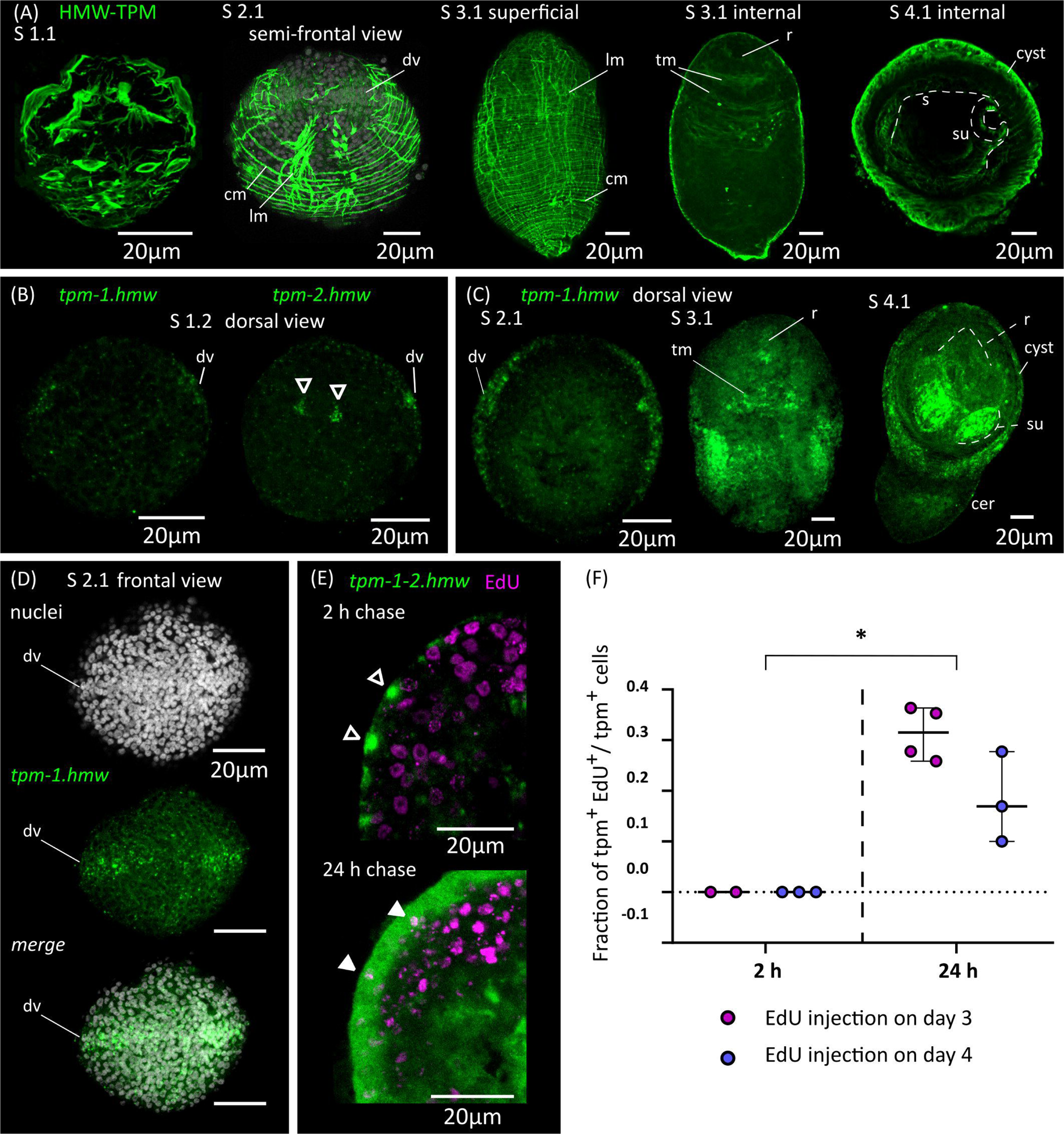
Development of the muscle system. **(A)** Detection of muscular tropomyosin isoforms by immunofluorescence at different developmental stages. Nuclear staining is included in greyscale for stage 2.1. For stage 3.1, both superficial planes and internal planes of confocal microscopy are shown. All pictures are from WMIHF except stage 4.1, which is from immunofluorescence on a cryosection. **(B)** Earliest detection of mRNAs for muscular tropomyosin isoforms *tpm-1.hmw* and *tpm-2.hmw* by WMISH at stage 1.2. Arrowheads indicate a pair of myocytons that probably correspond to the early asymmetric bundles of longitudinal muscle fibers. **(C)** WMISH of *tpm-1.hmw* at different developmental stages (dorsal views). **(D)** WMISH of *tpm-1.hmw* at stage 2.1 (frontal view). **(E)** Examples of larvae combining detection of EdU incorporation and expression of *tpm-1.hmw* and *tpm-2.hmw* by WMISH at different chase times after the injection of EdU to infected beetles. Open and filled arrowheads indicate *tpm1-2.hmw*^+^/EdU^-^ and *tpm1-2.hmw*^+^/EdU^+^ cells, respectively. **(F)** Quantification of the fraction of muscle cells (*tpm-1-2.hmw*^+^) that are labelled by EdU at different chase times after the injection of EdU to infected beetles. Each point represents the quantification of an individual larva (* p < 0.0025, Mann-Whitney test for pooled results of 3 and 4 d.p.i.). Abbreviations: cer, cercomer; cm, circular muscle; dv, band of cells at dorsoventral midline; lm, longitudinal muscle; r, rostellar primordium; s, scolex; su, sucker; tm, transverse muscle.

**Figure 5.**
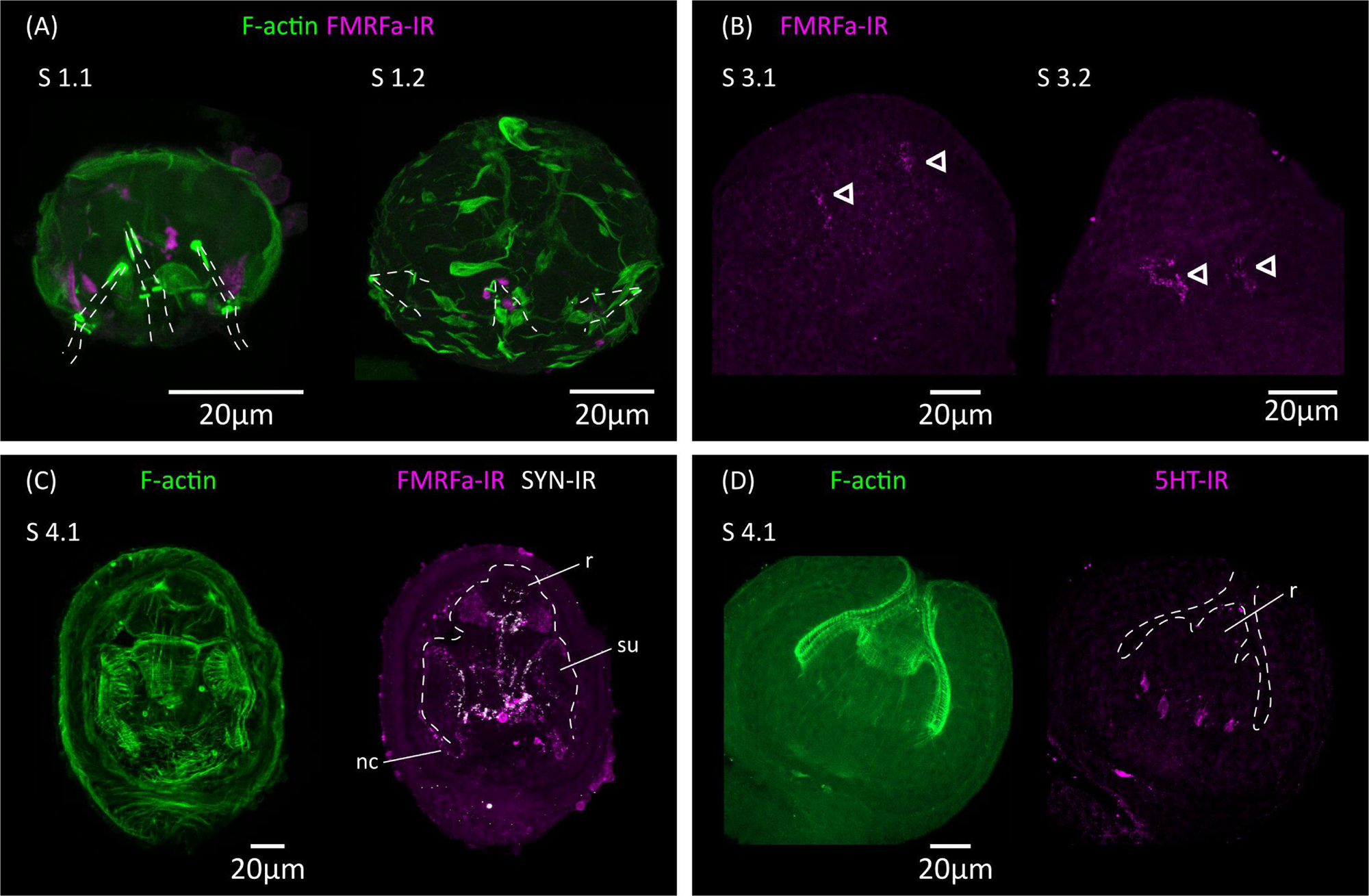
Analysis of the development of the nervous system by immunofluorescence. **(A)** FMRFa-IR in vestigial structures from the oncosphere during stages 1.1 and 1.2. Phalloidin staining of actin filaments (F-actin) labels the disorganized muscle fibers at these early stages. The positions of the larval hooks are indicated with dashed lines. **(B)** Earliest detection of FMRFa-IR in the developing scolex during stages 3.1 and 3.2. Arrowheads indicate early FMRFa-IR cells at the base of the rostellar primordium **(C)** FMRFa-IR, synapsin immunoreactivity (SYN-IR) and phalloidin staining of actin filaments during later scolex development (stage 4.1). **(D)** Earliest detection of 5-HT-IR occurs after scolex retraction (stage 4.1). Abbreviations: nc, nerve cord; r, developing rostellum; su, developing sucker.

In tapeworms, muscle fibers are only connected by thin cytoplasmic strands to their cell body containing the nucleus (myocyton) (Conn, 1993), and it is not usually possible to identify the myocytons by immunodetection of tropomyosins. Therefore, we performed WMISH with probes for muscular tropomyosin isoforms (*tpm-1.hmw* and *tpm-2.hmw*; Koziol et al., 2011) to identify myocitons during larval metamorphosis (Figure 4B, C, D). Both probes gave similar results, except that *tpm.hmw-1* was specifically absent from the developing cercomer. The earliest robust signal that we could detect was at stage 1.2, in cells forming a band at the lateral margin (marginal myocytons, located at the dorsoventral midline), and in the single pair of asymmetric myocytons associated with the early bundles of longitudinal muscle fibers (Figure 4B). Later, at stage 2.1 a second band of myocytons can be detected in the anterior end at the sagittal midline (*i*.*e*. perpendicular to the marginal myocytons), thus forming a characteristic cross when larvae are viewed frontally (Figure 4D). Strikingly, the position of the bands of myocytons can be distinguished by DAPI nuclear staining, with compactly distributed nuclei that have a characteristic zipper-like distribution (Figure 4D). At later stages (2.2 to 3.2), the marginal myocyton band becomes thicker, and more myocytons appear throughout the surface of the larvae, especially in the tissues that will become the cyst, behind the scolex. Additional myocytons appear among the inner cells (these are the myocytons of the transverse muscle fibers). Expression of muscular tropomyosins could be detected in the rostellar primordium at stage 3.2, but expression in the developing suckers only occurred after scolex withdrawal (stages 4.1 and 4.2, Figure 4C). At these stages, large numbers of myocytons were still present in the cyst tissues, and some were also detected in the growing cercomer. Altogether, these results expand those obtained by immunofluorescence, and indicate that muscle cell differentiation begins early during larval metamorphosis.

In order to detect the differentiation of germinative cells into muscle cells, we combined EdU chase experiments with WMISH detection of muscular tropomyosin isoforms (Figure 4E,F; for these experiments, we used a mixture of both *tpm1.hmw* and *tpm2.hmw* probes to detect all myocytons). When larvae were exposed to EdU at either 3 or 4 d.p.i., no colocalization was observed after 2 hours of chase between EdU labelling and expression of muscular tropomyosin isoforms, indicating that expression only begins after exiting the cell cycle (n=4-37 tropomyosin cells from 2-3 larvae for each time point, in which all positive cells were analyzed for each larva; identical results were observed in additional experiment using only either the *tpm1.hmw* or the *tpm2.hmw* probe). In contrast, extensive co-localization was observed after 24 hours of chase (31% of all myocytons were labelled in average after 24 hours when EdU was injected at 3 d.p.i, and 18% were labelled in average when EdU was injected at 4 d.p.i.; n=33-101 tropomyosin cells from 3-4 larvae for each time point). These results confirmed that extensive muscle cell differentiation is ongoing at 3 to 5 d.p.i. (stages 1.2 to 4.1), and demonstrated that the expression of muscle-specific effector genes begins 24 hours or less after cell cycle exit.

Finally, we analyzed the expression of homologs of *myoD* and *nkx-1.1*, which have been shown to be expressed in planarians in longitudinal muscle fiber cells and circular muscle fiber cells, respectively (Scimone et al., 2017) (Supplementary Figure 1). Furthermore, orthologs of *myoD* have been shown to be expressed and have myogenic activity in muscle progenitors and/or precursors in both vertebrates and invertebrates (Andrikou and Arnone, 2015). Expression of *myoD* during larval metamorphosis in *H. microstoma* was previously examined by Olson et al. (2018), and our results are similar to theirs, showing particularly strong expression in the future cyst tissues at stage 3.2, similar to the pattern of muscle tropomyosin isoforms at his stage. However, careful analysis showed that *myoD* expression was specifically absent in the band of marginal myocytons. On the other hand, *nkx-1.1* expression was detected from stage 3.1 onwards in the developing sucker primordia, with the strongest expression occurring after scolex withdrawal. Therefore, both genes are likely to be expressed in subsets of muscle cells and muscle cell precursors, but these may not be equivalent to those of planarians.

### *De novo* development of the nervous system of the cysticercoid

Ultrastructural analyses have identified a minimal nervous system in the oncosphere of many cestodes, including in the closely related *Hymenolepis nana*, which may comprise only two nerve cells that extend neurites towards the hook muscles (Fairweather and Threadgold, 1983; Swiderski et al., 2018). The oncosphere of *H. microstoma* has at least three nerve cells, which can be detected with antibodies generated against the neuropeptide FMRFamide and by their expression of synapsin (Preza, 2021). We analyzed the development of the nervous system during larval metamorphosis by detecting FMRFamide immunoreactivity (FMRFa-IR), 5-hydroxytryptamine (serotonin) immunoreactivity (5-HT-IR), and synapsin protein expression (Figure 5).

During the first 3 d.p.i. (stages 1.1 to 2.1), FMRFa-IR was detected in spots associated to the vestigial oncosphere hooks at the larval posterior, which later disappeared (Figure 5A). These structures are likely the degenerating remains of the nervous system of the oncosphere. In contrast, FMRFa-IR cells in the developing scolex appeared at 5 d.p.i in stages 3.1 to 3.2 as a pair of weakly positive cells at the base of the rostellar primordium (Figure 5B). Extensive FMRFa-IR and synapsin could only be detected after scolex withdrawal, starting at stages 4.1, and included the developing brain commissure and longitudinal nerve cords (Figure 5C).

On the other hand, 5-HT-IR was only detected in the scolex after scolex withdrawal, first as four positive cells associated to the base of the developing rostellum (Figure 5D), and later more extensively in positive cells in the scolex and in the developing longitudinal nerve cords. These results strongly indicate that the nervous system of the cysticercoid develops independently from that of the oncosphere, given the spatial and temporal separation in their development.

In addition, we analyzed by WMISH the expression of markers of different neurotransmitter systems present in *H. microstoma*, which are also expressed in the nervous system of the adult worm (Preza et al., 2018): prohormone convertase 2 (*pc2*) as a marker of peptidergic cells; choline acetyltransferase (*chat*) as a marker of cholinergic cells; vesicular glutamate transporter (*vglut*) as a marker of glutamatergic cells; and tryptophan hydroxylase (*tph*) as a marker of serotonergic cells (Figure 6). Expression of *pc2*, *chat* and *vglut* could be detected from 4 d.p.i. onwards (at stage 2.1), in a few cells at the anterior end, and the number of positive cells increased at stages 2.2 to 3.2. Expression at these early stages was associated with the anterior-most structures of the nervous system, including the developing rostellar ganglia, cerebral ganglia and transverse commissure. Development continued after scolex withdrawal, with expanded expression domains in the scolex, and with the appearance of the longitudinal nerve cords. On the other hand, *tph* expression began later at 5 d.p.i. (stage 3.2, Figure 6D), in few cells in the scolex, and the total number of positive cells was always small, which correlates with the small number of serotonergic cells found in the adult worm (Preza et al., 2018).

**Figure 6.**
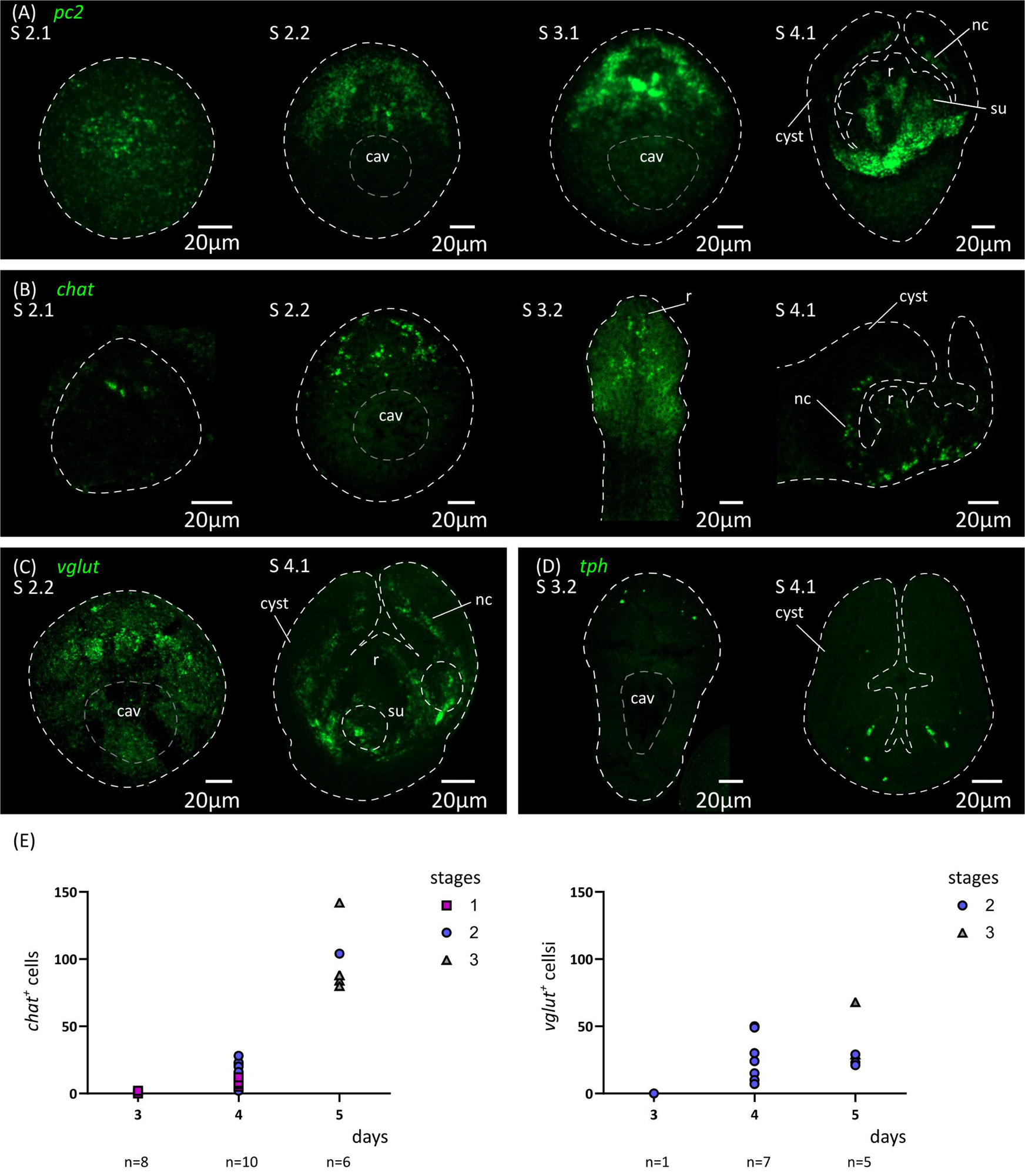
Analysis of the development of the nervous system by *in situ* hybridization of markers of neurotransmission. Expression patterns of *pc2* **(A)**, *chat* **(B)**, *vglut* **(C)** and *tph* **(D)** during larval metamorphosis. **(E)** Quantification of the total number of *chat*^+^ and *vglut*^+^ cells at 3 to 5 d.p.i. Abbreviations: cav, central cavity; nc, nerve cord; su, sucker primordium.

Thus, the initial expression of markers of neurotransmission at the mRNA level preceded the appearance of FMRFa-IR, 5-HT-IR and synapsin protein expression by at least 24 hours, demonstrating that cell differentiation in the nervous system is already underway at stage 2.1. This suggests that neural progenitors should already be present at even earlier stages. We analyzed by WMISH the expression of homologs of genes coding for transcription factors that are related to neurogenesis and neural differentiation in animals, including *soxB* and the basic helix-loop-helix (bHLH) family members *coe* and *neuroD* (Dubois and Vincent, 2001; Cowles et al., 2013; Hartenstein and Stollewerk, 2015; Baker and Brown, 2018; Tutukova et al., 2021). All of these genes had expression patterns that were associated with the developing nervous system (Figure 7). Both *soxB* and *coe* were detected in relatively small numbers of cells in the developing nervous system from stage 2.1 onwards, and showed little incorporation of EdU after two hours of labelling (8% of *soxB*^+^ cells incorporated EdU, n=189 cells from 2 larvae; 2% of *coe*^+^ cells incorporated EdU, n=179 cells from 3 larvae), suggesting that most of these cells correspond to post-mitotic neural precursors. In contrast, *neuroD* could be detected at least from stage 1.2 throughout the anterior hemisphere of the larvae (we did not examine its expression at earlier time points), in large proliferative cells: approximately 80% of all *neuroD*^+^ cells were labelled by EdU after 2 hours, and *neuroD*^+^ cells correspond to 14% of all EdU^+^ proliferative cells in early larvae (n=110 *neuroD*^+^ cells and 836 EdU^+^ cells examined from 6 larvae; Figure 8A). Expression of *neuroD* was also observed in mitotic cells. During later development, *neuroD*^+^ cells were always loosely distributed around regions of ongoing neural development. After scolex withdrawal, *neuroD* expression was mostly turned off in the scolex but continued around the developing nerve cords (Figure 7A). Therefore, *neuroD* appears to be expressed early during neurogenesis, marking a sizable subpopulation of all proliferating germinative cells.

**Figure 7.**
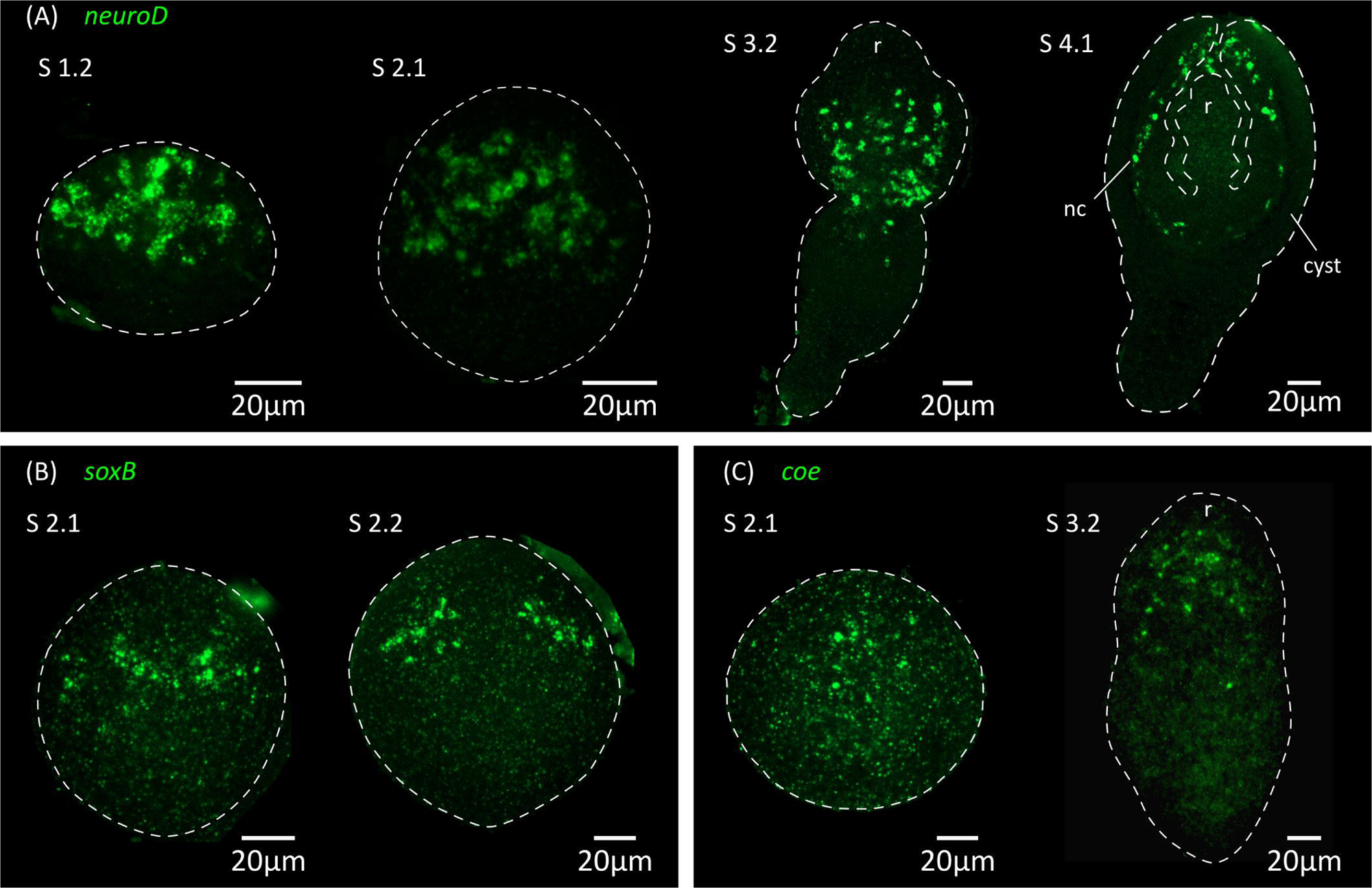
Expression of conserved genes related to neurogenesis and neural differentiation. Expression patterns of *neuroD* **(A)**, *soxB* **(B)** and *coe* **(C)** during larval metamorphosis. Abbreviations: nc, nerve cord; r, rostellar primordium.

**Figure 8.**
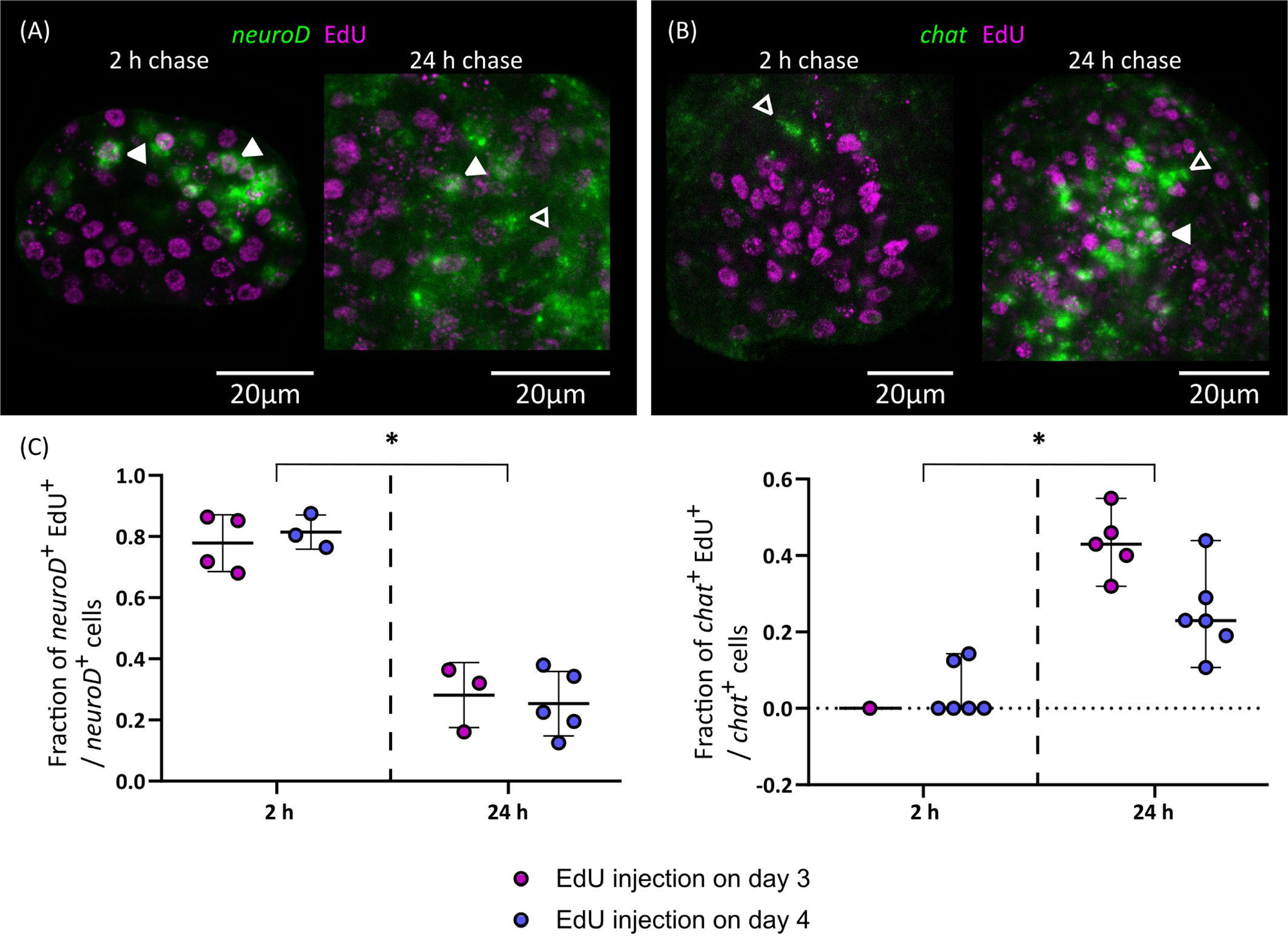
Differentiation of nerve cells during larval metamorphosis. Examples of EdU labelling of *neuroD*^+^ cells **(A)** and *chat*^+^ cells **(B)** after 2 or 24 h of chase (open and filled arrowheads indicate EdU^-^ and EdU^+^ cells, respectively), and quantification of the fraction of *neuroD*^+^ **(C)** or *chat*^+^ **(D)** cells that are labelled by EdU at different chase times. Each point represents the quantification of an individual larva (* p < 0.001, Mann-Whitney test for pooled results of 3 and 4 d.p.i.).

Finally, we traced the differentiation of nerve cells by means of EdU chase assays. We focused on *chat* as a marker of neural differentiation because it was expressed in an intermediate number of spread-out cells, which made the identification of the nuclei of positive cells non-ambiguous. We injected EdU in infected beetles at 3 and 4 d.p.i., and analyzed EdU labeling in *chat*^+^ cells after 2 or 24 hours. Either no or very few *chat*^+^ cells could be detected after two hours (as these corresponded to larvae at stages 1.2 to 2.1) and most *chat*^+^ cells in these larvae were EdU negative (Figures 8B and 8C). However, for two larvae, a single weakly positive *chat*^+^ cell was found for each. Analysis of larvae at later stages labelled for two hours with EdU also showed no co-localization between *chat* and EdU (data not shown). Therefore, *chat* expression appears to be turned on after cell cycle exit in most cases. In comparison, 24 hours after EdU injection there was extensive co-localization of *chat* and EdU (43% and 25% of all *chat*^+^ cells are EdU^+^ after 24 hours, when injection occurred 3 d.p.i or 4 d.p.i., respectively; Figures 8B and 8C). Thus, neurogenesis must be already underway at stages 1.2 to 2.1, and neural differentiation begins within 24 hours of cell cycle exit. In contrast, the proportion of *neuroD*^+^ EdU^+^ cells decreased after 24 hours, suggesting that these cells are turning off *neuroD* expression as they differentiate during this period, and/or that they proliferate extensively, resulting in the dilution of the EdU label (Figures 8A and 8C).

## Discussion

In this work, we have characterized cell proliferation and differentiation of new tissues during the larval metamorphosis of a model tapeworm. This developmental transition in tapeworms has largely been regarded as a “black box”, since tracing the development and remodelling of the larval tissues is complicated due to the extremely small size of the larvae and their cells, the complex histology of cestodes, and the lack of specific markers. Thus, most of what was previously known depended on spatial and temporal snapshots from studies by electron microscopy. Here, we took advantage of the development of specific molecular markers and a robust *in vivo* system for metabolic labelling with a thymidine analogue, allowing us to obtain a global view of these processes (summarized in Figure 1).

The first stages of larval metamorphosis are characterized by exponential growth, as has also been seen during the cysticercoid to adult transition (Loehr and Mead, 1980). In both cases, set-aside germinative cells are thought to be quiescent before infection, and to become quickly activated during the infection of the new host. During the first stages of larval metamorphosis in *H. microstoma*, we could detect the differentiation of tegumental cells and muscle cells. The early differentiation of new tegument cytons has also been described during the larval metamorphosis of *Echinococcus multilocularis* (Sakamoto and Sugimura, 1970), and may be essential for the profound changes that the tegument undergoes during the early metamorphosis. It is likely that these changes are important for the survival of the parasite in the new host (Holcman and Heath, 1997). The early differentiation of muscle cells could be related to their multiple functions in tapeworms. On the one hand, muscle activity is required for scolex withdrawal (Caley, 1974), which occurs after only 5 to 6 days of metamorphosis. On the other hand, myocitons are known sources of signalling molecules regulating development in tapeworms and other flatworms (Witchley et al., 2013; Koziol et al., 2016; Díaz-Soria et al., 2020; Wendt et al., 2020), and are the main source of extracellular matrix components as has also been shown in planarians (Conn, 1993; Witchley et al., 2013; Cote et al., 2019). The distribution of myocytons during early metamorphosis is very peculiar, as they are initially concentrated on a lateral marginal band (at the dorsoventral midline), which is followed by a second band at the sagittal midline. A similar pattern was observed during the early development of the *E. multilocularis* protoscolex (Koziol et al., 2014). It is possible, given their position and their time of appearance, that these are the myocytons of the first circular muscle fibers that appear, although we have not been able to confirm their cytoplasmic continuity. As is common in invertebrates, *myoD* was not expressed in all somatic muscles (Tixier et al., 2010; Andrikou and Arnone, 2015; Brunet et al., 2016), and it was specifically absent from the marginal myocytons of *H. microstoma*. Because in planarians *myoD* is expressed in longitudinal muscles (Scimone et al., 2017), this would support their tentative classification as circular muscle cells. However, although *nkx-1.1* is expressed in circular muscle cells in planarians (Scimone et al., 2017), it was restricted to the developing suckers in *H. microstoma*. Therefore, it may not be possible to homologize muscle cell types across platyhelminthes based on their expression of conserved myogenic transcription factors.

The development of the nervous system occurs with a clear antero-posterior gradient, similar to that found in *E. multilocularis* protoscoleces (Koziol et al., 2013). Differentiation of the nervous system begins relatively late during metamorphosis, and scolex withdrawal occurs at a stage in which the nervous system is still under development (and at which synapsin labelling is still undetectable). Therefore, it is possible that the muscle contractions responsible for scolex withdrawal are not controlled or modulated by the nervous system. Strikingly, the nervous system of the cysticercoid is formed post-embryonically *de novo*, independently of the nervous system of the oncosphere. It is reminiscent of the development of the nervous system during the embryonic development in other ectolecithal flatworms (*i*.*e*. with alecithal oocytes and specialized yolk cells carrying the vitellum), where the nervous system does not arise from a specialized region of the ectoderm, but from progenitors found among a large internal mass of proliferating cells (Hartenstein and Stollewerk, 2015; Monjo and Romero, 2015). Our results suggest that *neuroD*^+^ germinative cells may correspond to a specialized lineage of neural precursors during larval metamorphosis. Surprisingly, *neuroD* appears to be an early expressed gene during neurogenesis in *H. microstoma*, in contrast to its more common expression in late neuronal precursors during differentiation in other animals (Monjo and Romero, 2015; Sur et al., 2020; Deryckere et al., 2021; Tutukova et al., 2021), with the exception of the annelid *Platynereis dumerilii* (Simionato et al., 2008). Small numbers of proliferating neoblasts have also been shown to express a *neuroD* homolog in adult planarians (Scimone et al., 2014). On the other hand, *soxB* had a restricted expression in *H. microstoma* larvae and was expressed later in what appears to be a subset of mostly post-mitotic precursors. This is also surprising, since *soxB* homologs usually are expressed during the early specification of the neuroectoderm in animal embryos (Hartenstein and Stollewerk, 2015), although later roles during neuronal differentiation have also been described for particular *soxB* homologs (Guth and Wegner, 2008; Phochanukul and Russell, 2010; Monjo and Romero, 2015; Vidal et al., 2015). Furthermore, the ortholog of *soxB* in *E. multilocularis* is expressed in most proliferating germinative cells in the metacestode germinative layer (Cheng et al., 2017), which are unlikely to be neural precursors given the minimal nervous system present in this tissue (Koziol et al., 2013). Finally, tapeworms appear to lack many conserved regulators of neural development, such as *neurogenin* and *elav* (Montagne and Koziol, unpublished data), hinting at a highly modified neurogenic program. It would be interesting to determine if these atypical patterns of gene expression also occur during neural development in other life stages, including embryonic development and during the post-embryonic remodelling of the nervous system of the adult. More generally, our gene expression results demonstrate the molecular heterogeneity of tapeworm germinative cells. Stem cell heterogeneity was also hinted at by ultrastructural details during the early larval metamorphosis in *E. multilocularis* (Sakamoto and Sugimura, 1970), and shown from differential gene expression in fully developed metacestodes of this species (Koziol et al., 2014). In adults of the related tapeworm *Hymenolepis diminuta*, different genes were also shown to be expressed either exclusively in some proliferating germinative cells, or in subsets of germinative cells and in differentiated tissues, suggesting the existence of lineage-restricted germinative cells (Rozario et al., 2019).

Altogether, our results support the hypothesis that larval metamorphosis corresponds to a form of ′maximal indirect development′ (Peterson et al., 1997; Koziol, 2017), in which a new body plan is generated *de novo* from stem cells that were set-aside during embryogenesis. We have found no clear evidence of any differentiated cells from the oncosphere that are retained in the cysticercoid after metamorphosis. Instead, new cells of the nervous system, muscular system and tegument are generated from the differentiation of germinative cells. Although we did not analyze the development of the excretory system, this must also arise *de novo* during metamorphosis as it is absent in the oncosphere of *Hymenolepis* spp. (Ubelaker, 1980). *H. microstoma* appears as an ideal model to study the mechanisms regulating cell proliferation and differentiation during larval metamorphosis, given its simple maintenance and fast development. This work provides a blueprint of the early stages of development, and the *in vivo* system could be complemented in the future by functional analyses in *in vitro* culture (Seidel, 1975). We expect that the insights gained with this model species may guide future studies in human and animal pathogens.

### Ethics statement

The protocol for the maintenance of *Hymenolepis microstoma* in mice has been approved by Comisión Honoraria de Experimentación Animal, Uruguay (protocol number 10190000025215)

### Author contributions

UK conceived the project. JM and UK designed the experiments. JM, MP and UK performed the experiments and analyzed the data. UK wrote the initial draft of the manuscript. All authors contributed to the article and approved the submitted version.

## Acknowledgments

The authors would like to acknowledge the collaboration of Jenny Saldaña, Laboratorio de Experimentación Animal, Facultad de Química, Universidad de la República, for the maintenance of the life cycle of *Hymenolepis microstoma* in the laboratory. The authors would also like to acknowledge the help of Anita Asienberg and Laura Montes de Oca (Departamento de Ecología y Biología Evolutiva, Instituto de Investigaciones Biológicas Clemente Estable, Montevideo, Uruguay) with *T. molitor* rearing and Beatriz Goñi (Facultad de Ciencias, Universidad de la República) for her advice regarding the manipulation of *T. molitor* beetles. The authors gratefully acknowledge the Advanced Bioimaging Unit at the Institut Pasteur Montevideo, and Gabriela Casanova, Magela Rodao and Gaby Martínez at the Electron Microscopy Unit, Facultad de Ciencias, Universidad de la República, for their support and assistance in the present work.

## Figure legends

**Supplementary Figure 1.** Expression of *myoD* **(A)** and *nkx-1.1* **(B)** homologs during larval metamorphosis. Abbreviations: dv, lateral band of marginal myocitons; su, sucker primordia / developing sucker.

**Supplementary Table 1**. List of primers used in this work.

