## Supplementary figures and images for "Stem cell proliferation and differentiation during larval metamorphosis of the model tapeworm *Hymenolepis microstoma*"

### Supplementary Figure 1

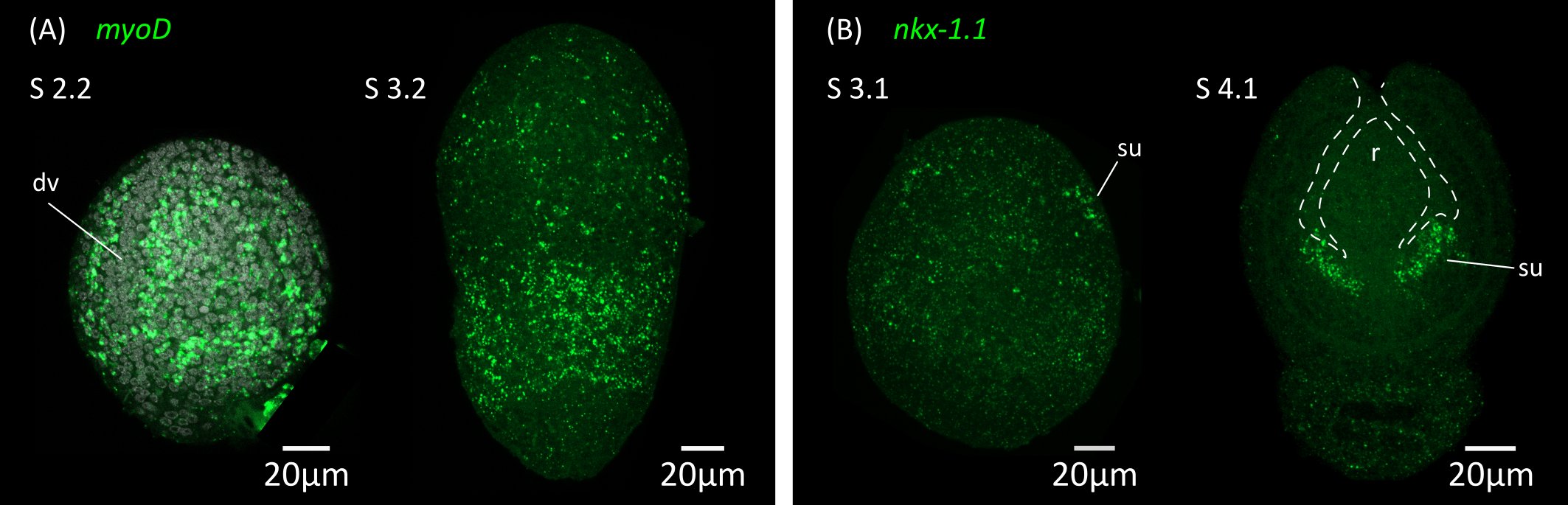
